# Long-term salinity reveals genotype-specific transcriptional reprogramming in eggplant

**DOI:** 10.64898/2026.01.13.699247

**Authors:** Matteo Martina, Cristina Morabito, Andrea Moglia, Anna Maria Milani, Lorenzo Barchi, Alberto Acquadro, Cinzia Comino, Francesca Secchi, Ezio Portis

## Abstract

Salinity severely limits eggplant productivity, yet the transcriptional bases of tolerance to prolonged salt exposure remain incompletely understood. Here, we analyzed long-term salinity responses in two contrasting eggplant (Solanum melongena L.) genotypes from the G2P-SOL core collection, focusing on genotype-dependent transcriptional regulation under chronic stress. Plants were exposed to 200 mM NaCl for 23 days at the reproductive stage, and transcriptome profiling was performed at the end of the stress period. Physiological assessment and high-throughput phenotyping confirmed a strong divergence in water status and plant architecture between genotypes under salinity, providing a reference framework for transcriptomic interpretation. RNA-seq analysis revealed marked genotype-specific differences in transcriptional responses. While both genotypes activated a conserved salt-stress program involving redox homeostasis, proteostasis and growth repression, the tolerant genotype displayed a substantially broader and more coordinated transcriptional reprogramming. This response involved large-scale modulation of pathways related to translation and RNA metabolism, hormone signaling crosstalk, membrane transport, cell wall remodeling and oxidative stress management, together with the selective repression of growth- and signaling-related functions. In contrast, the sensitive genotype showed a more limited response dominated by defense- and damage-associated transcripts. Overall, these results indicate that long-term salt tolerance in eggplant is associated with genotype-specific transcriptional reprogramming superimposed on a shared basal stress response. This work highlights regulatory pathways and candidate genes potentially relevant for breeding strategies targeting salt resilience.

## Introduction

Salinity is one of the major abiotic constraints limiting crop productivity in many irrigated regions, and its incidence is expected to increase further under climate change scenarios [1, 2]. Eggplant (*Solanum melongena* L.) is widely cultivated in Mediterranean and subtropical environments, where soil and water salinization are becoming increasingly common [2, 3]. Although eggplant is often classified as moderately tolerant to salinity compared with other Solanaceae, yield reductions and alterations in fruit quality are frequently reported under saline conditions [4–9]. Physiological and biochemical investigations have shown that salt stress in eggplant is associated with reductions in photosynthetic performance and plant water status, accompanied by alterations in ion homeostasis, antioxidant activity and osmolyte accumulation. Importantly, tolerant genotypes tend to maintain higher leaf water potential, improved redox balance and more effective stress acclimation mechanisms compared with sensitive ones [4, 5, 7, 10]. Despite these advances, the physiological traits contributing to salt tolerance often show quantitative and genotype-specific patterns, complicating their direct translation into breeding strategies.

At the molecular level, transcriptomic studies in eggplant have begun to identify salt-responsive genes and pathways, including components related to hormone signaling, redox regulation, cell wall remodeling and primary metabolism [11–13]. However, most available transcriptomic analyses have focused on early stress responses, specific tissues (e.g. roots or seedlings) or single genotypes, providing only a partial view of the regulatory strategies deployed under salinity. Consequently, the transcriptional programs underlying genotype-dependent tolerance to prolonged salinity, particularly at the reproductive stage, remain poorly characterized.

In recent years, the G2P-SOL project (https://www.g2p-sol.eu/) has assembled and characterized an eggplant core collection capturing the broad genetic and geographic diversity of the crop [14, 15]. This collection has been phenotyped for multiple traits (agronomical, quality and metabolic traits, and to both biotic and abiotic stresses - [16]), providing a powerful resource to identify contrasting genotypes and to link phenotypes with genetic and genomic information.

The development of high-throughput phenotyping platforms now allows non-destructive, repeated measurements of plant architecture and canopy development over time [17, 18]. 3D imaging systems, such as PlantEyeⓇ, can quantify traits like projected leaf area, canopy height, voxel-based volume and digital biomass, providing a detailed picture of how plants grow and adjust under stress. When combined with classical physiological measurements, such as leaf water potential, these tools offer a powerful framework to characterize salt responses at the whole-plant level [19, 20].

In this study, two contrasting lines for tolerance to soil salinity were selected within the G2P-SOL eggplant core collection, both physiological and molecular approaches were used to characterize eggplant response to long-term salinity at the reproductive stage. After almost four weeks of the salt experiment, one line exhibited high water status, whereas the second line showed a strong decline in leaf water potential. 3D imaging and RNA-seq were performed at the end of the stress period, aiming to i) provide a picture of the canopy state of two lines under stress, ii) characterize the global transcriptomic responses to salt stress in the two contrasting lines, and iii) identify genotype-specific pathways and molecular mechanisms associated with superior tolerance.

## Methods

### Plant material and pre-selection of contrasting genotypes

Two contrasting eggplant genotypes were selected from the G2P-SOL core collection (http://www.g2p-sol.eu/). The Indian landrace ‘TS00870’ (GPE036890) was chosen because it had previously been characterized as tolerant to salinity, whereas the Spanish cultivar ‘Berenjena del terreno’ (GPE022290) had shown clear sensitivity to salt stress in earlier G2P-SOL project trials [21]. Seeds were sown and grown under controlled environmental conditions in a growth chamber. The chamber conditions were maintained at 25°C/18°C (day/night), with a photoperiod of 16 h light/8 h dark and relative humidity of approximately 60–70%. At 10 leaves stage, seedlings were transferred to 5L pots containing a standardized soil mixture and transferred to the greenhouse.

### Salinity treatment

Five months-old plants were splitted in two irrigation regimes: control (CTR-Hoagland solution - [22]) and salt treatment (STR - Hoagland solution added with NaCl, final concentration 200 mM). The salinity treatment was imposed at the reproductive stage and maintained for 23 days. Control and salt-treated plants of each genotype were arranged in a randomized design on the glasshouse bench. Three biological replicates per genotype × treatment combination were used for leaf water potential measurements and RNA-seq sampling at the end of the trial.

### Leaf water potential measurements

Leaf water potential (Ψ_leaf) was assessed at the end of the 23-day salinity period. Measurements were taken at midday on fully expanded transpiring leaves using a Scholander-type pressure chamber, following standard procedures. For each genotype and treatment, three individual plants were selected and one leaf per plant was measured. These data were used to quantify the impact of salinity on plant water status and to compare the responses of the salt sensitive line GPE022290 and the tolerant line GPE036890.

### 3D imaging and pot weight monitoring

Plants of the two eggplant genotypes under control and salt treatment were monitored using a PlantEye 3D multispectral scanner (Phenospex, Heerlen, The Netherlands) mounted on a linear rail above the canopy, provided with an automated weighing system (Droughtspotter, Phenospex), which recorded pot weight in parallel with imaging. For each genotype × treatment combination, a single representative plant was selected for 3D imaging, with the aim of obtaining high-resolution, time-resolved reconstructions of whole-plant architecture and canopy structure rather than population-level statistical estimates. Plants were scanned once per day over the last four days of salt treatment (days 20–23), when plants were at the reproductive stage. For each scan, PlantEyeⓇ software was used to extract canopy structural and optical traits, including 3D leaf area (3DLA), mean surface angle (leaf inclination - SA) and the green leaf index (GLI). A stressed/control index was calculated separately for each genotype as the ratio between the value of the salt-treated plant and that of the corresponding control plant. Time courses of these indices over the four imaging days were used for graphical representation and to compute the area under the curve (AUC) for each genotype and trait in R. Accordingly, 3D imaging data were interpreted as descriptive and illustrative of genotype-specific architectural responses and were not used for inferential statistical testing. In addition, 3D point clouds exported from PlantEyeⓇ were imported into CloudCompare (www.cloudcompare.org) to reconstruct and visualize the plant architecture, providing a qualitative representation of whole-plant geometry and canopy structure under control and salt conditions.

### RNA extraction and sequencing

For transcriptome profiling, fully expanded leaves were sampled at the end of the salinity treatment from control and salt-stressed plants of the two eggplant genotypes. Sampling was carried out on the same plants and at the same time point used for leaf water potential measurements. For each genotype and treatment, three biological replicates were collected, each consisting of a pooled leaves sample from an individual plant, which was immediately frozen in liquid nitrogen and stored at −80 °C until processing. Total RNA was extracted from approximately 100 mg of powdered leaf tissue using a column-based plant RNA purification kit (Spectrum™ Plant Total RNA Kit, Sigma-Aldrich), followed by DNase treatment to remove residual genomic DNA. RNA concentration and purity were assessed spectrophotometrically, and RNA integrity was verified by agarose gel electrophoresis and capillary electrophoresis. Stranded mRNA libraries were prepared from poly(A)-enriched RNA according to standard Illumina protocols and sequenced as paired-end reads (2 × 150 bp) on an Illumina platform at IGAtech (Udine, Italy).

### Read processing, transcript quantification and differential expression analysis

Raw sequencing reads were subjected to quality control and adapter trimming to remove low-quality bases and residual adapter sequences (*fastp -* [23]). Cleaned reads were then pseudo-aligned to the *S. melongena* reference transcriptome corresponding to the ‘67/3’ eggplant line genome assembly (SMEL v5;[16]) using Salmon in quasi-mapping mode with default parameters [24]. Transcript-level abundances were estimated in terms of counts and Transcripts Per Million (TPM), and gene-level counts were subsequently imported in R [25] using the *tximport* package [26]. All downstream statistical analyses were performed in R with the DESeq2 package [27]. The Deseq2 design formula included the main effects of genotype and stress and their interaction, allowing to distinguish constitutive differences between genotypes from treatment-specific transcriptional responses. Size-factor normalization and dispersion estimation were carried out following the standard DESeq2 workflow, and Wald tests were used to obtain log₂ fold changes and adjusted p-values for each gene. From this model, within-genotype contrasts comparing salt-treated and control plants in each line, as well as between-genotype contrasts comparing the two genotypes under control and salinity conditions, were extracted. The interaction term between genotype and stress was also inspected to identify genes showing genotype-specific responses to salinity. *p-values* were adjusted for multiple testing using the Benjamini–Hochberg procedure [28], and genes were considered differentially expressed when the adjusted *p-value* was below 0.1. For heatmap visualization, a subset of highly regulated DEGs was selected based on both statistical significance and effect size (|log₂ fold change| > 1) and further filtered to retain genes showing consistent expression differences across biological replicates. For exploratory visualization, normalized counts were transformed by applying a *log₂(count + 1)* transformation after size-factor normalization. These transformed values were used for principal component analysis and for the construction of expression heatmaps of the most strongly regulated genes, generated with the *pheatmap* package [29].

### Gene set enrichment analysis (GSEA)

To investigate transcriptional changes at the pathway level, Enriched GO categories were visualised using the enrichplot package (https://github.com/YuLab-SMU/enrichplot) in R. For each contrast of interest, all genes with non-missing log₂ fold change were ranked according to the DESeq2 log₂ fold change, with positive values indicating higher expression in the second level of the contrast. These ranked gene lists were used as input for the gseGO function, employing a dedicated S. melongena annotation package and SMEL5 gene identifiers as keys. Gene Ontology enrichment was carried out separately for the Biological Process, Molecular Function and Cellular Component ontologies, using a minimum gene set size of 5 and a maximum of 500 genes. Significance of enrichment scores was assessed using a simple permutation scheme with 20,000 permutations, and p-values were adjusted for multiple testing using the Benjamini–Hochberg procedure. GO terms with an adjusted p-value ≤ 0.05 were considered significantly enriched. Enriched GO categories were visualized using the enrichplot package. Ridgeplots were generated to display, for each GO term, the distribution of ranked log₂ fold changes of core enriched genes, with color scales encoding either the normalized enrichment score or the adjusted p-value. In addition, mirrored dot plots were constructed to compare enrichment patterns between the two genotypes by jointly displaying, for each GO term, its presence in GPE022290 and/or GPE036890, and the size of the corresponding core gene sets. This representation allowed us to distinguish terms that were uniquely enriched in the sensitive line, uniquely enriched in the tolerant line, or shared between them.

## Results

### Leaf water potential reveals contrasting salt tolerance between two eggplant genotypes

Leaf water potential was measured at the end of the salt treatment in control and NaCl-treated plants of both lines (Figure 1). Under control conditions, the two genotypes showed comparable leaf water potential values (mean ± SD) of −0.51 ± 0.12 MPa in GPE022290 and −0.36 ± 0.09 MPa in GPE036890. After 23 days of salinity, leaf water potential drops in both lines, but the decline was much stronger in GPE022290 (−1.78 ± 0.10 MPa) than in GPE036890 (−0.59 ± 0.29 MPa). Stressed plants of line GPE036890 maintained leaf water potentials similar to those of the controls, whereas line GPE022290 exhibited markedly more negative values, revealing contrasting water status under salt stress between the two genotypes.

**Figure 1.**
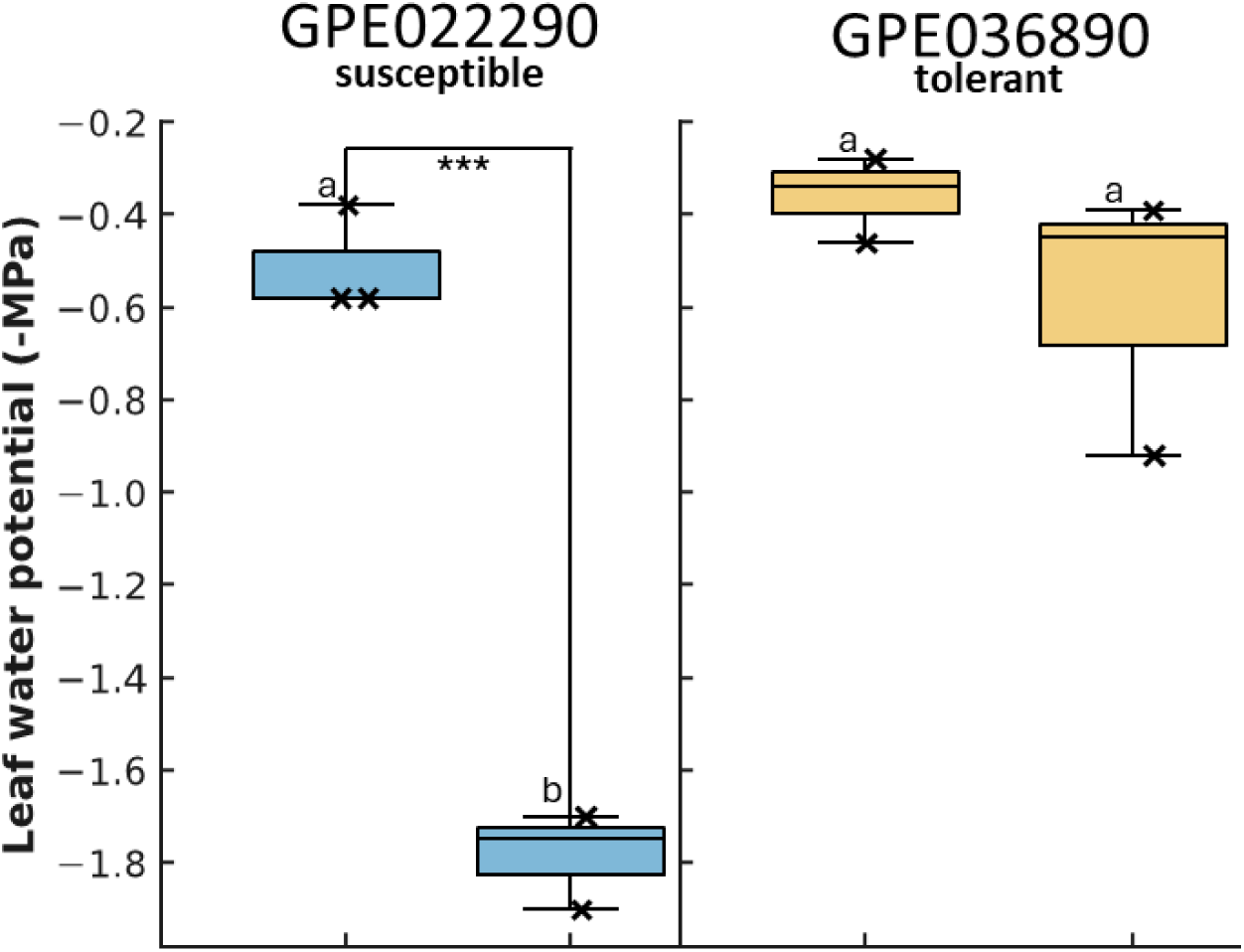
Leaf water potential (Ψ_leaf) of the sensitive genotype (GPE022290) and the tolerant genotype (GPE036890) under control (CTR) and salt-stress (STR) conditions after 23 days of NaCl treatment. Boxes show median and interquartile range. Different letters above the boxes indicate significant differences among genotype × treatment combinations (P < 0.05), whereas asterisks denote significant differences between control and salt-treated plants within each genotype (* P < 0.05; ** P < 0.01; *** P < 0.001).

### Canopy architecture dynamics under prolonged salinity stress assessed by 3D imaging

Representative 3D reconstructions of the plants illustrate how prolonged salinity reshapes canopy architecture in the two genotypes. Under control conditions (Figure 2 A,C), canopies appear tall and compact, with leaves filling most of the reconstruction volume, whereas salt-treated plants (Figure 2 B,D) show more open and discontinuous canopies and a marked increase in leaf drooping and twisting. These qualitative differences in plant architecture motivated a quantitative analysis of 3D-derived traits (e.g. leaf area, surface angle and GLI) over time to compare the canopy responses of the salt sensitive (Figure 2 A,B) and tolerant lines (Figure 2 C, D).

**Figure 2.**
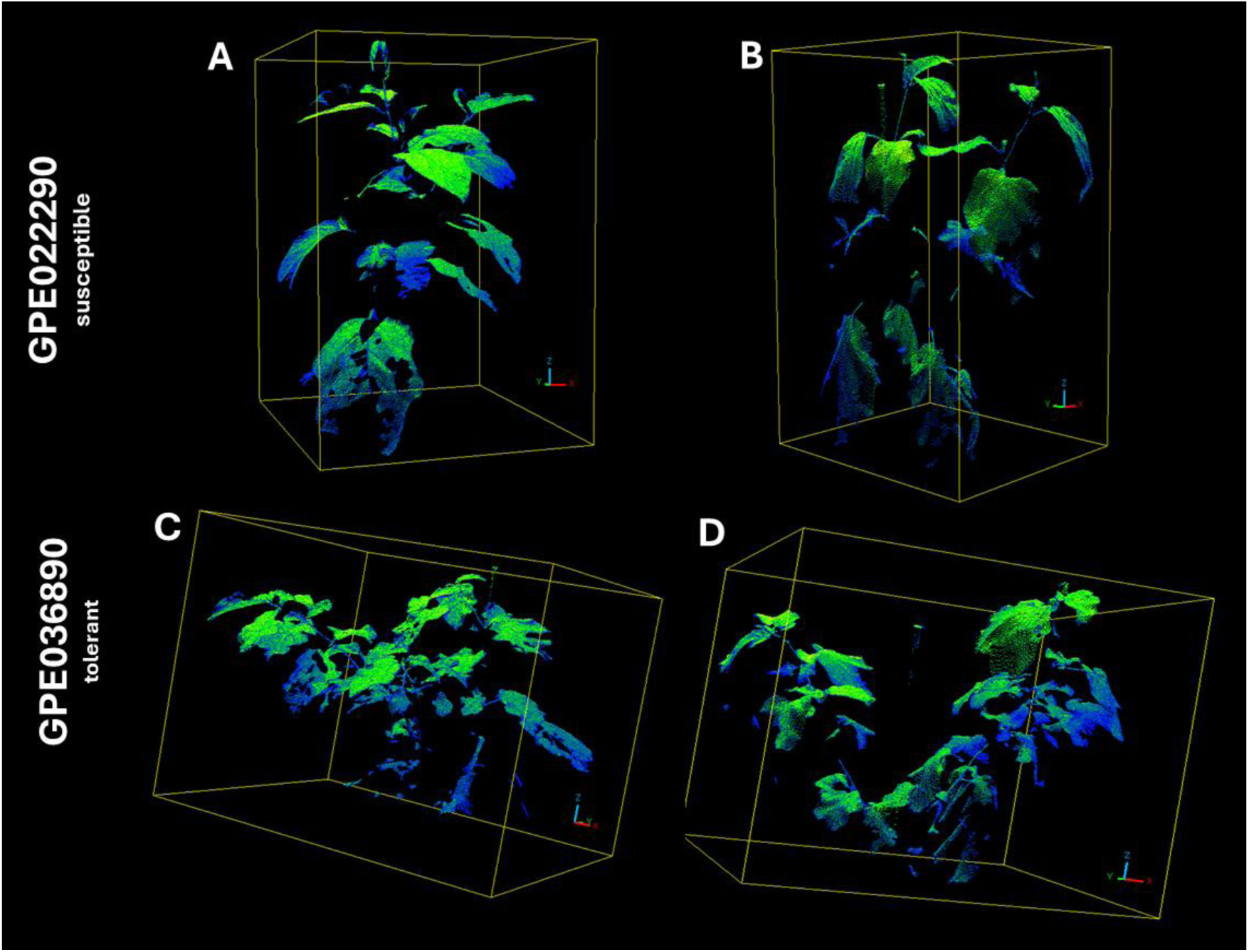
Representative 3D point clouds acquired with the PlantEye scanner for GPE022290 (A, B) and GPE036890 (C, D) after 23 days of treatment. Panels A and C show plants grown under control conditions (CTR), whereas panels B and D show plants exposed to NaCl (STR). The color gradient (blue to green) reflects canopy reflectance/greenness.

Stressed/control indices were monitored for 3D leaf area, surface angle, GLI and pot weight during the last three days of the experiment (Figure 3). In both genotypes, the 3D leaf area index decreased over time, with values ranging from 1.25 to 1.08 in line GPE036890 and from 0.63 to 0.52 in line GPE022290 (Fig. 3A). Surface angle indices also progressively declined in both genotypes between days 0 and 3, with values spanning 0.85–0.77 (-10%) in GPE036890 and 0.85–0.71 (-16,5%) in GPE022290 (Fig. 3B). GLI indices remained close to 1 in line GPE036890 (1.02–0.98) and slightly below 1 in line GPE022290 (0.86–0.80) over the same period (Fig. 3C). Pot weight indices were consistently greater than 1 throughout the same period for both genotypes, ranging from 1.12 to 1.24 in GPE036890 and from 1.22 to 1.33 in GPE022290 (Fig. 3D).

**Figure 3.**
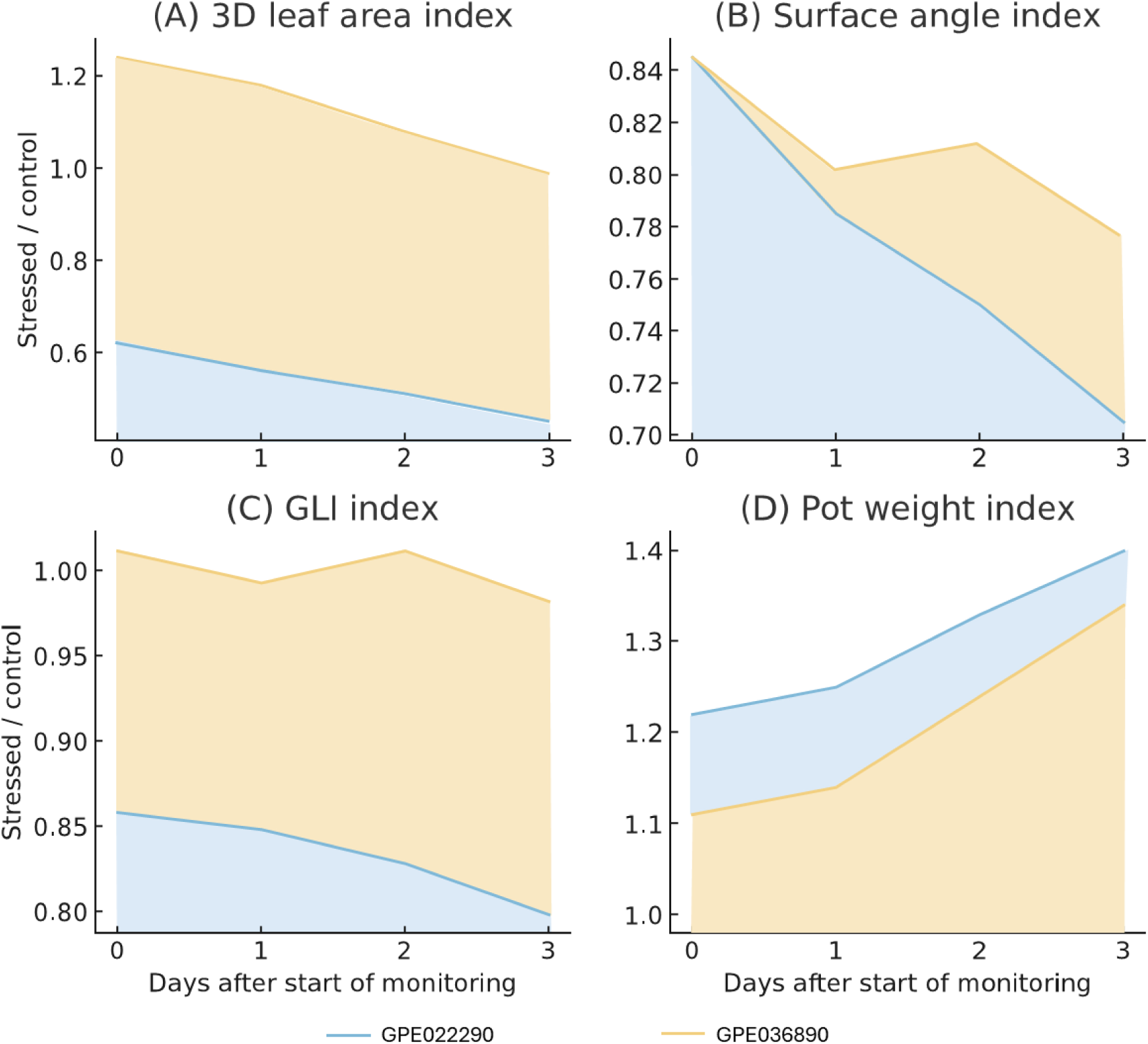
Temporal dynamics of stress indices (stressed/control ratios) for GPE022290 (blue) and GPE036890 (yellow) across the last three days of NaCl treatment. (A) 3D leaf area index, (B) surface angle index, and (C) greenness index (GLI) were derived from PlantEye 3D imaging, whereas (D) pot-weight index reflects cumulative changes in water content level. Shaded areas represent the area under the curve (AUC), highlighting the cumulative temporal response of each genotype over the observation period.

The late response of the two genotypes was summarized by calculating the area under the curve (AUC) of stressed/control indices for 3D leaf area, surface angle, GLI and pot weight. The AUC of the 3D leaf area index was 3.37 for line GPE036890 and 1.63 for line GPE022290. For the surface angle index, the AUC values were 2.30 for line GPE022290 and 2.43 for line GPE036890. The AUC of the GLI index was 2.50 for line GPE022290 and 3.00 for line GPE036890. The AUC of the pot weight index was 3.89 for line GPE022290 and 3.62 for line GPE036890.

### Global transcriptomic responses to salinity in sensitive and tolerant lines

To compare the global transcriptional responses to salinity in the susceptible and tolerant lines, we performed RNA sequencing on control and salt-treated plants sampled at the end of the stress period, using three independent biological replicates per genotype × treatment combination, in accordance with current RNA-seq experimental standards. Raw reads (∼40 million paired end reads per sample) were subjected to quality filtering and aligned to the eggplant reference transcriptome (SMEL v5; Gaccione et al., 2025). Transcript-level abundances were estimated, and gene-level differential expression analyses were performed using DESeq2.

PCA clearly separated the two genotypes along the first principal component, reflecting their strong constitutive divergence in gene expression profiles (Figure 4A). The second component discriminated against control and salt-treated samples within each genotype. Only 560 up-regulated and 347 down-regulated genes were detected in salt-treated susceptible plants compared with their controls (Figure 4B-C), while the tolerant one showed a broader transcriptional reprogramming under salinity, with 1428 up-regulated and 1306 down-regulated genes (Figure 4B-D).

**Figure 4.**
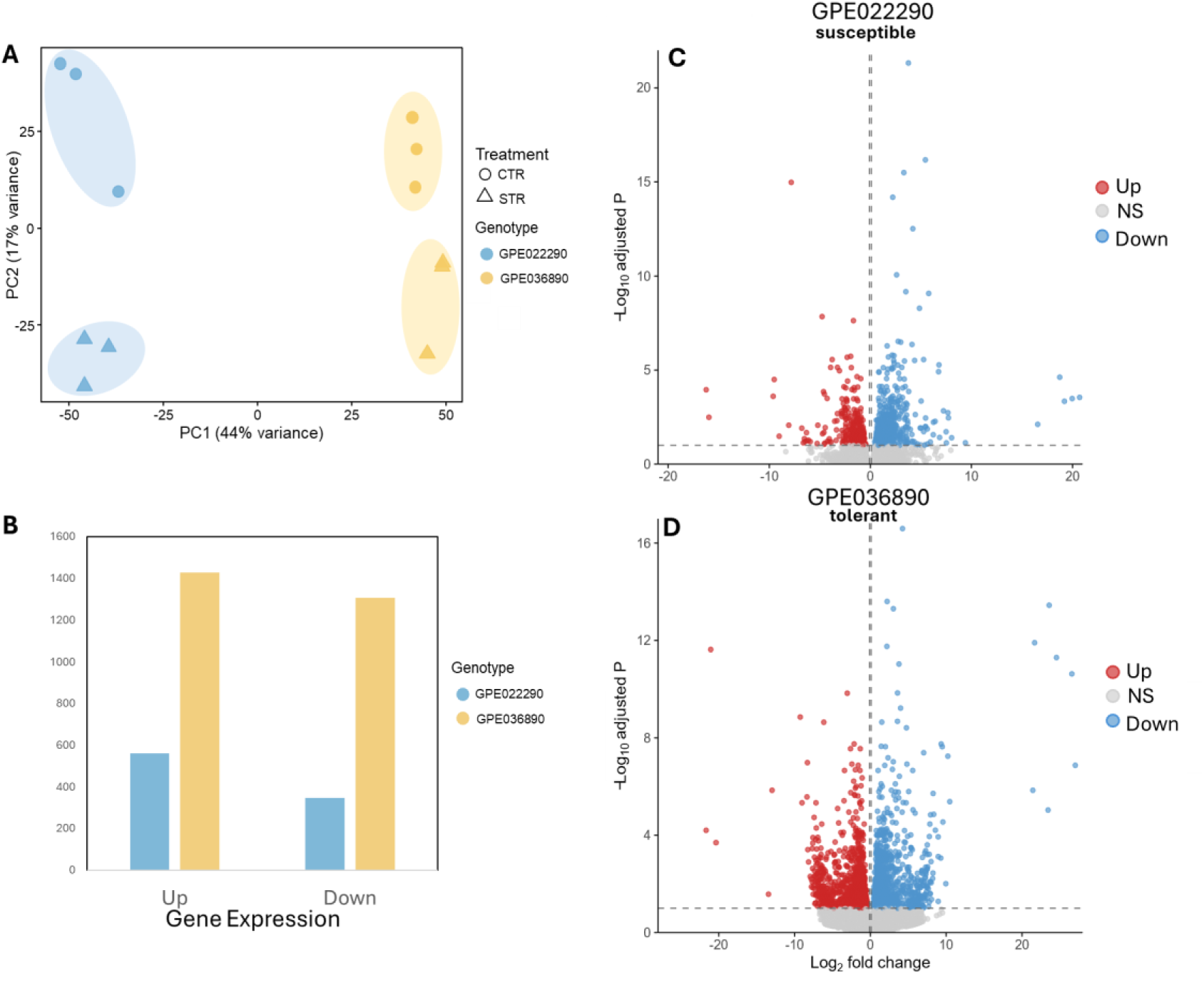
Global transcriptional response to salinity in contrasting eggplant genotypes GPE022290 and GPE036890. (A) Principal component analysis (PCA) based on variance-stabilised counts from RNA-seq data of GPE022290 (blue) and GPE036890 (yellow) under control (circles) and salt-stress (triangles) conditions; (B) Number of differentially expressed genes (DEGs) identified in each genotype, separated into up- and down-regulated genes; (C) Volcano plot showing the distribution of differentially expressed genes (DEGs) in the sensitive genotype GPE022290 and (D) the tolerant genotype GPE036890 after salt treatment. Blue and red dots indicate significantly up- and down-regulated genes, respectively. The horizontal dashed line marks the adjusted P-value threshold.

The most strongly differentially expressed genes were investigated by visualizing their expression patterns in a heatmap. Heatmaps were constructed using a subset of highly regulated DEGs selected based on both effect size and adjusted p-value and further filtered to ensure coherent expression patterns across biological replicates. This representation allowed us to assess the reproducibility of the transcriptional response and to evaluate sample clustering according to treatment and genotype (Figure 5).

**Figure 5.**
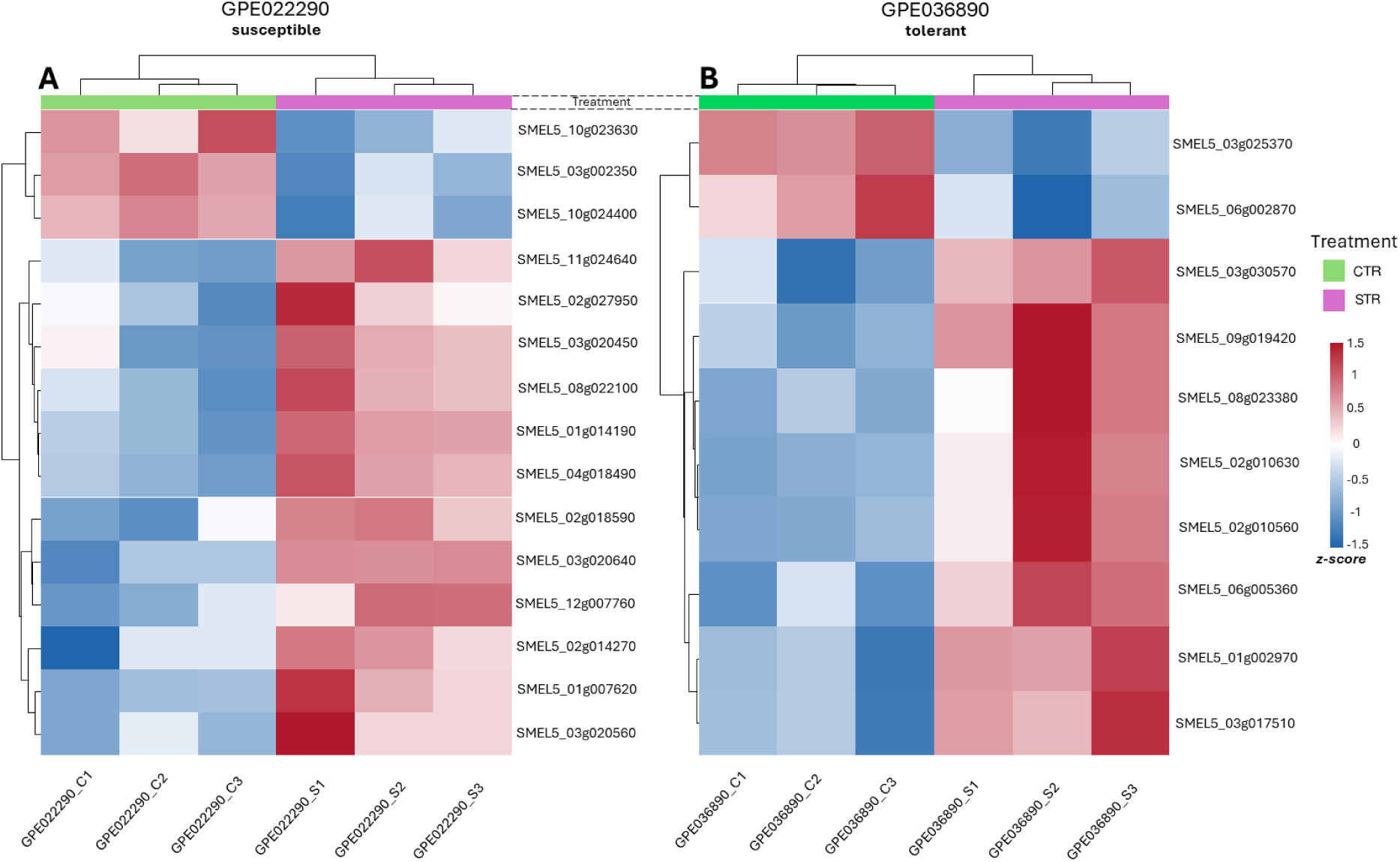
Heatmaps of genotype-specific transcriptional responses to salt stress in contrasting eggplant lines. Expression heatmaps of the top differentially expressed genes (DEGs) under control (CTR) and salt stress (STR) conditions in (A) the sensitive genotype GPE022290 and (B) the tolerant genotype GPE036890. Colors represent normalized expression values (Z-scores), with red indicating upregulation and blue downregulation relative to the mean. Each column corresponds to an individual biological replicate.

In the susceptible line, the expression profiles of the top-ranked differentially expressed genes resulted in a clear separation between control and salt-treated samples, with all stress replicates clustering together and showing a coherent expression pattern (Figure 5A). Most of these transcripts were strongly induced by salinity, displaying low expression in control plants and high expression under salt stress. The gene set appeared largely enriched in defense- and stress-related functions, including pathogenesis-related proteins (SMEL5_01g014190, SMEL5_08g022100, SMEL5_02g018590), β-1,3-glucanase (SMEL5_03g002350), multiple protease inhibitors (SMEL5_01g007620, SMEL5_03g020560, SMEL5_03g020640, SMEL5_03g020450), enzymes involved in oxidative and specialized metabolism (SMEL5_02g014270, SMEL5_02g027950, SMEL5_04g018490, SMEL5_10g023630), as well as the auxin-related transporter (SMEL5_10g024400) and the plastidial aminotransferase (SMEL5_12g007760). Together, these genes allowed to define a compact, stress-induced expression group of genes that characterizes the response of line GPE022290 to prolonged salinity (Supplementary Table 1).

In the tolerant line, the heatmap of the most strongly differentially expressed genes also showed a clear separation between control and salt-treated samples, with replicates clustering together and displaying a consistent expression pattern (Figure 5B). Most of these transcripts were up-regulated under salinity and include genes associated with signaling and regulation (SMEL5_01g002970), apocarotenoid metabolism (SMEL5_02g010560, SMEL5_02g010630, SMEL5_09g019420), cell wall–related or nutrient-responsive proteins (SMEL5_03g030570, SMEL5_06g002870), redox and metal homeostasis (SMEL5_06g005360, SMEL5_03g025370), and defense-associated protein (SMEL5_08g023380). Together, these DEGs define a coordinated transcriptional set of genes that is activated by salt in line GPE036890 (Supplementary Table 1).

To move from gene-level patterns to pathway-level responses, we next performed Gene Set Enrichment Analysis (GSEA) using ranked log₂ fold changes for each contrast. Enriched Gene Ontology terms in the Biological Process, Cellular Component and Molecular Function categories were then summarized using dot plots, which provide a compact view of the number of core enriched genes in each genotype under control and salt conditions. GO enrichment based on GSEA highlighted marked differences between the two lines at the level of molecular functions, biological processes and cellular components (Figure 6). Within the molecular function (MF) category (Fig. 7A), most enriched terms were specific to line GPE036890 and were supported by large core gene sets, including oxidoreductase activities, various UDP-dependent glycosyltransferases acting on benzoic acid, nicotinate and salicylic acid, terpene and sesquiterpene synthase activities, and structural/structural-constituent functions of the ribosome. In contrast, line GPE022290 contributed only a few MF terms, mainly related to cytoskeletal and microtubule motor activity, with very limited overlap between the two genotypes. A similar pattern was observed for biological process terms (Fig7B), where GPE036890 showed enrichment for multiple pathways involved in hormone and isoprenoid metabolism, secondary cell wall biogenesis and RNA processing, whereas the susceptible line accounted for only a small subset of categories. In the cellular component (CC) category (Fig.7C), the two lines showed almost complementary signatures. Enriched CC terms in GPE022290 were largely associated with ribosomes and cytoskeletal structures, whereas the tolerant line was characterized by terms related to the cell wall and extracellular/apoplastic compartments.

**Figure 6.**
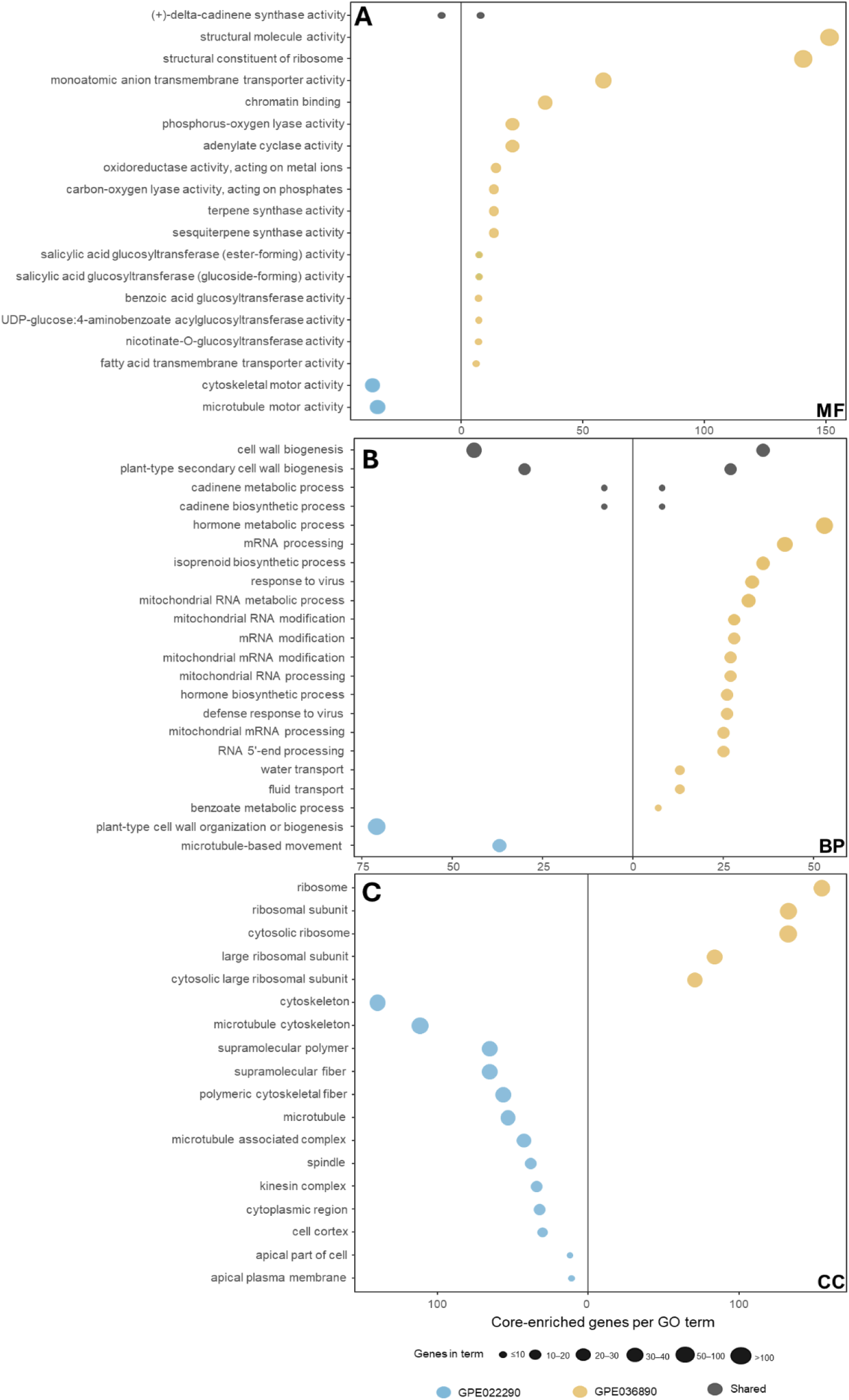
Gene set enrichment analysis (GSEA) of salt-induced transcriptional changes in contrasting eggplant genotypes. Mirrored dot plots showing significantly enriched Gene Ontology (GO) terms for the biological process (A), cellular component (B) and molecular function (C) domains in the sensitive genotype GPE022290 (blue, left) and the tolerant genotype GPE036890 (yellow, right). The x-axis and dot size represent the number of genes per GO term.

### Genotype-specific transcriptional signatures associated with salt tolerance in genotype GPE036890

Comparison of salt-induced DEGs between the two genotypes revealed a markedly broader transcriptional reprogramming in the tolerant line GPE036890 (Figure 7). Among upregulated genes, 1.153 (67.3%) were specific to GPE036890, whereas only 285 (16.6%) were unique to the sensitive GPE022290 and 275 (16.1%) were shared by both genotypes. A similar pattern was observed for downregulated genes: 1.134 (78.2%) were specifically repressed in GPE036890, compared with 145 (10.0%) unique to GPE022290 and 172 (11.9%) common to both lines, suggesting that salinity tolerance in GPE036890 might be associated with an extensive, line-specific transcriptional reprogramming rather than with a small set of shared core responses.

**Figure 7.**
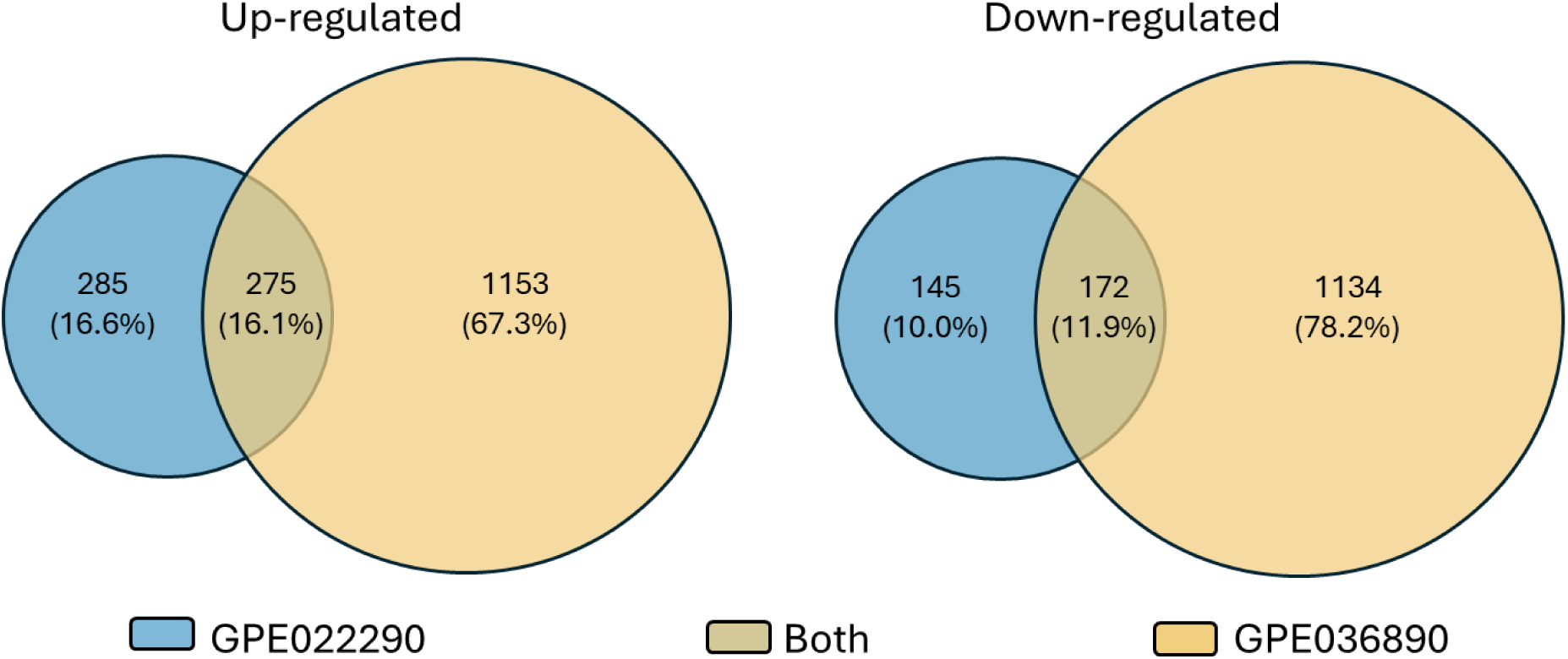
Venn diagrams showing the number and proportion of significantly upregulated (top) and downregulated (bottom) genes shared or unique between the sensitive genotype GPE022290 (blue) and the tolerant genotype GPE036890 (yellow). Percentages indicate the relative contribution of each category to the total number of DEGs.

GSEA revealed that only a limited subset was specifically enriched in the sensitive line GPE022290 (Figure 7). Comparing the two lines under stress conditions, GPE036890 showed strong positive enrichment for categories BP related to protein synthesis and RNA metabolism linked to translation, such as rRNA processing, tRNA metabolic process and ribosome biogenesis, while multiple terms associated with mitochondrial and nuclear RNA processing (including mitochondrial RNA modification/processing and RNA 5’-end processing) were negatively enriched (Figure 8). Consistently, the CC analysis highlighted positive enrichment of ribosome and cytosolic ribosome, together with plastidial membrane systems (thylakoid- and photosynthetic-membrane–related terms) specifically in GPE036890. In the MF families, GPE036890 showed strong positive enrichment of structural constituents of ribosome and structural molecule activity, whereas catalytic activities such as adenylate cyclase and phosphorus–oxygen lyase were significantly depleted in the tolerant line. Together, the GSEA profiles indicate that under salinity, large, coordinated shifts in ribosomal, organellar and redox-related gene sets, combined with a selective downregulation of specific RNA-processing and signaling functions, are a hallmark of the tolerant genotype, whereas enrichment in the sensitive line is restricted to a narrower set of cytoskeleton- and cell-wall-related functions.

**Figure 8.**
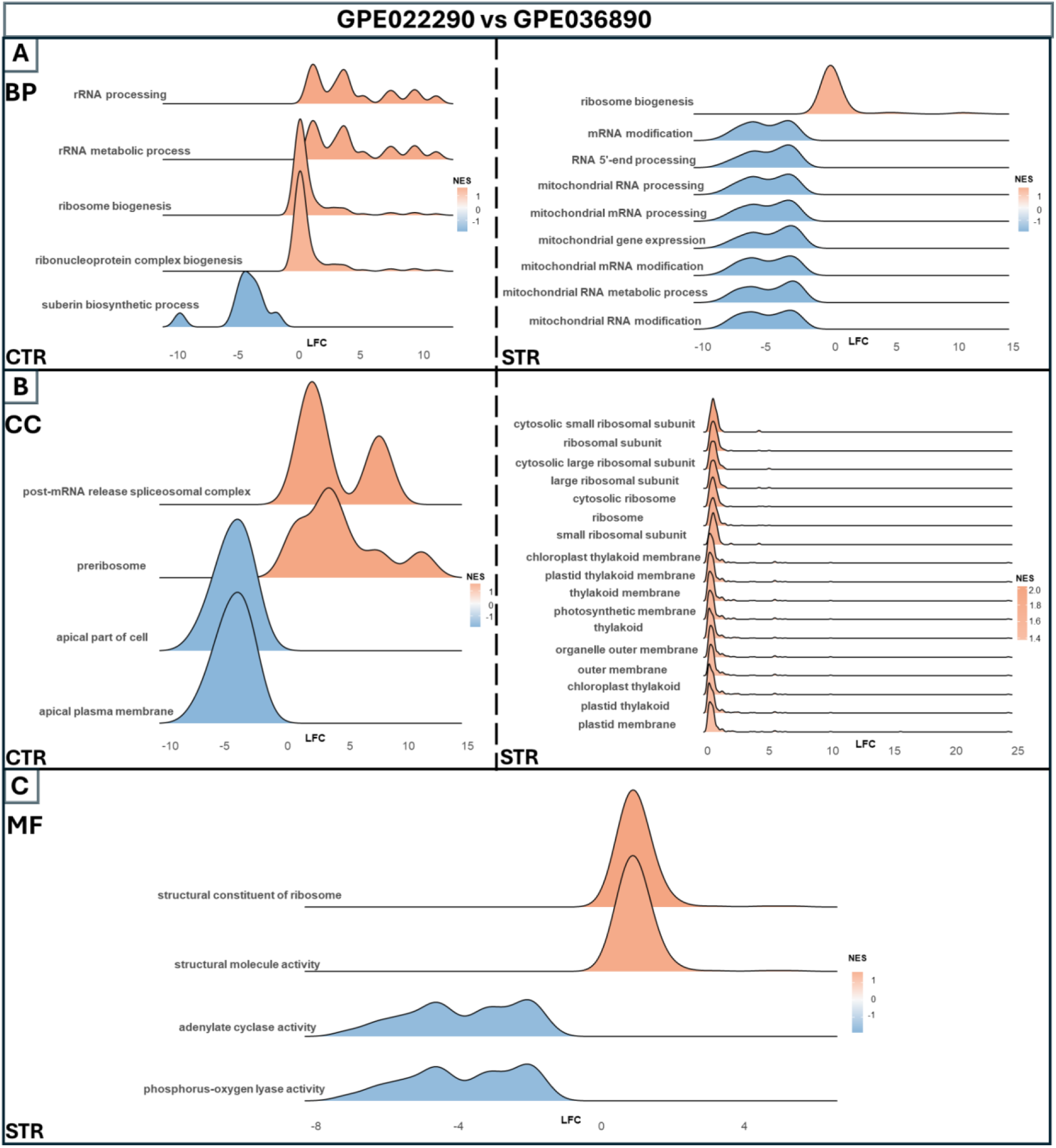
Comparative gene set enrichment analysis (GSEA) between eggplant genotypes under control and salt-stress conditions. Ridge plots showing significantly enriched GO terms in the biological process (A), cellular component (B) and molecular function (C) categories for the tolerant genotype GPE036890 and the sensitive genotype GPE022290 under control (left panels) and salt-stress (right panels) conditions. The x-axis represents log₂ fold change (LFC), and the distribution shape reflects the gene-level contribution to each term. Curves are filled with NES value, ranging from negative (blue) to positive (orange) values.

## Discussion

The present study focuses on the transcriptomic dissection of prolonged salinity responses in two contrasting eggplant genotypes selected from the G2P-SOL core collection. Physiological measurements and 3D canopy phenotyping were used to characterize stress progression and to define comparable stress states across genotypes, guiding the interpretation of transcriptomic sampling.

### Leaf water potential and canopy traits as indicators of salt tolerance

Leaf water potential provided a clear separation between the two genotypes. After 23 days of NaCl treatment, GPE022290 dropped the Ψ_leaf_ values close to −2.0 MPa, while GPE036890 remained near control levels (around −0.5 MPa). These values are consistent with limited osmotic adjustment and/or impaired water uptake in the sensitive line. In contrast, the tolerant genotype appears to buffer the external salinity, presumably through more efficient osmotic adjustment and ion compartmentation, thereby maintaining turgor despite high NaCl watering conditions.

3D imaging and pot-weight measurements allowed to link these differences to canopy behavior. Stressed/control indices for 3D leaf area and GLI suggest that GPE036890 experiences only modest reductions in projected canopy size and green leaf fraction, whereas GPE022290 shows a pronounced loss of leaf area and greenness. Moreover, pot-weight indices above 1 in both genotypes indicate a reduction in evapotranspiration under salt, with slightly higher values in GPE022290 pointing to a stronger impairment of water use.

### Global transcriptomic responses

Principal component analysis showed that most variance in gene expression reflects constitutive differences between the two genotypes, with the salt treatment superimposed on this pre-existing divergence. In GPE022290, relatively few genes passed the significance thresholds and most transcripts remained close to log₂FC ≈ 0, suggesting a limited capacity to remodel the transcriptome after prolonged salt exposure. The emerging DEGs are mainly associated with protease inhibitors, chitinases, pathogenesis-related proteins, and β-1,3-glucanases, together with oxidative and specialized metabolism enzymes and a limited set of regulatory genes. This profile, dominated by canonical PR hydrolases and defense regulators, closely resembles a generic defense-/damage-associated program, typically observed upon severe tissue injury or pathogen attack, rather than the classic osmotic adjustment and ion homeostasis machinery mobilized under salt tolerance (e.g. ion transporters of the SOS/HKT/NHX families - [30, 31] - , compatible solute biosynthesis - [32, 33] - , LEA/dehydrin proteins - [34–36] - and ABA-dependent signaling - [37–39]), further supporting the view that this genotype mainly does not operate an efficient osmotic adjustment strategy.

By contrast, GPE036890 displays a much broader pattern of transcriptional change, with large gene sets being both induced and repressed. Strongly up-regulated/down-regulated genes encompass signaling components, enzymes of apocarotenoid and specialized metabolism, cell wall– and nutrient-responsive proteins, and factors involved in redox and metal homeostasis. Rather than being confined to a single defense-like mechanism, the transcriptional changes in GPE036890 span metabolism, transport, structural adjustment and stress signaling, pointing to a coordinated reprogramming of growth, resource allocation and cellular homeostasis under salinity. This type of multi-layered adjustment is typically associated with effective salt tolerance strategies, in which osmotic/ionic homeostasis, ROS management and maintenance of photosynthetic tissues are jointly regulated, rather than sacrificed in favor of short-term defense [40, 41]. Consistently, the tolerant line maintains a more favorable physiological status (Figure 1,2), preserving canopy structure and greenness under salinity (Figure 2,3), in line with a genuine acclimation/tolerance response rather than with a predominantly damage-driven syndrome.

### A conserved salt-stress program shared by tolerant and sensitive genotypes

Despite the strong genotype-specific component, our data highlights a coherent set of pathways consistently modulated by salinity in both lines (Figure 6 - Supplementary Table S1). These shared changes define a conserved transcriptional salt-stress program that likely represents the basal adjustment required to cope with osmotic and ionic stress, irrespective of the final tolerance phenotype.

Together with a common rebalancing of hormone signaling (Figure 6B), involving the upregulation of ethylene biosynthetic genes [42], strigolactone signaling modulators [43, 44], and oxylipin/JA-related steps [45, 46], both genotypes up-regulate canonical stress-protection mechanisms, including LEA/dehydrins, HSP chaperones, universal stress and senescence-associated proteins, together with ubiquitin-dependent protein quality-control components (Figure 6A), suggesting enhanced proteostasis, protein quality control and membrane/protein protection as a common response to prolonged salinity in eggplant. In parallel, both lines strongly upregulated redox and ROS-management genes (Figure 6A), consistent with the expected oxidative burden under salt stress, and indicated that both genotypes invest in detoxification and fine-tuning of cellular redox poise [47–54]. The regulation of secondary-metabolism-related genes (Figure 6C) centered on phenylpropanoids and terpenoids indicates a redirection of carbon flow towards the synthesis of antioxidants [55–58] and incorporation of phenolic units into structural polymers such as lignin and suberin [59, 60]. The expression of cell wall–related proteins suggest a tendency toward the reinforcement and reconfiguration of the cell wall, impacting on porosity, mechanical properties and ion fluxes, and limiting mechanical and oxidative damage under salinity [61].

Genes commonly downregulated by salt define a complementary group associated with growth- and resource-demanding functions (Supplementary Table S1). This set includes regulators of cell expansion and development like auxin response factors [62], brassinosteroid and peptide signaling components [63, 64], and cell cycle-related regulators [65], pointing to a coordinated downshift of growth-promoting programs. In addition, salt stress remodel transport and photosynthesis-related functions, with selective repression of specific transporters and antenna components alongside induction of putative repair/protective modules and thylakoid function, and carbon allocation to primary metabolism and cell-wall biosynthesis, consistent with a reprioritization of carbon use, water transport capacity and membrane traffic away from growth and towards maintenance [66–71].

Overall, the convergence between shared GO categories and commonly regulated genes suggests that both genotypes activate a conserved salt-stress program centered on reinforcement of proteostasis and ROS detoxification, remodeling of secondary metabolism and the cell surface, extensive hormonal/signaling adjustments and concurrent repression of growth, photosynthesis and transport functions. This basal program is therefore unlikely to explain the contrasting salt tolerance by its presence or absence, but rather provides a shared framework on which genotype-specific quantitative differences are required to shape the final phenotype.

### Tolerant-specific transcriptional reprogramming in GPE036890

To dissect tolerance-linked regulation under prolonged salinity, we focused on DEGs uniquely responsive to stress in the tolerant line GPE036890 (Figure 7), revealing a coordinated reallocation of transcriptional investment consistent with long-term acclimation to osmotic and ionic stress. Rather than amplifying a single canonical pathway, the tolerant line remodels hormonal and signaling circuits, transport capacity, redox and proteostasis networks, and cell-surface architecture, while attenuating growth-promoting processes. Notably, several of the most strongly contrasting tolerant-specific genes do not belong to functional categories that are globally enriched in the direct genotype contrast under stress, indicating that tolerance is not solely driven by broad shifts in a limited number of GO classes but also relies on the selective engagement of key regulatory and effector genes embedded within more widely distributed functional backgrounds. All tolerant-specific DEGs discussed below, together with functional annotations, genomic coordinates, and line-specific log2 fold changes, are reported in Table 1.

A prominent axis of tolerant-specific reprogramming involves auxin signaling and homeostasis. This was reflected by the coherent enrichment of early auxin-responsive components, including SAUR71-like and SAUR68-like growth regulators, GH3.1-like auxin-conjugating enzymes, and Aux/IAA4-like repressors, together with regulators of auxin transport and distribution such as PIN-LIKES 7 and BIG. In parallel, the mild repression of ARF2A suggests a fine-tuning of auxin-dependent transcription rather than a generalized attenuation of the pathway. Overall, the coordinated modulation of multiple auxin-related functional categories (Table 1-Figure 8) points to an active buffering of auxin pools and responsiveness, consistent with current models in which auxin integrates growth–stress trade-offs and supports developmental plasticity under chronic salinity ([72]- Figure 8B). Auxin-associated changes were embedded within a broader hormone-related reprogramming. Functional categories linked to ethylene signaling were selectively over-represented, as indicated by the modulation of several ethylene-responsive transcription factors (ERF014, CRF4 and ERF003-like), together with brassinosteroid-responsive components of the EXORDIUM family. This pattern is consistent with extensive auxin–ethylene–brassinosteroid crosstalk governing growth restraint, cell expansion and stress adjustment [73]. Additional enrichment of categories related to gibberellin and cytokinin signaling further supports a multilayered hormonal coordination rather than the dominance of a single regulatory axis. In contrast, functional categories associated with canonical ABA biosynthesis and stress signaling [63], apart from the induction of abscisic stress-ripening protein and LEA14-A. This functional profile suggests that, at this advanced stage of salt exposure, acclimation relies primarily on protective and osmoprotective mechanisms, with limited engagement of an ABA-centered transcriptional program, pointing instead to predominantly ABA-independent strategies.

Several tolerant-specific modules in GPE036890 converge on core processes known to underpin acclimation to prolonged salinity. Functional categories associated with membrane transport and solute homeostasis were prominently represented, as reflected by the induction of a polyol transporter 5-like, the anion channel SLAH2-like, the chloride channel CLC-b, and multiple ABC transporters. The coordinated enrichment of transport-related functions points to a reinforced control of solute fluxes and membrane transport, a key requirement under chronic osmotic and ionic stress to stabilize cellular water relations and mitigate toxic ion accumulation ( [74–77] - Figure 6B). In parallel, categories linked to redox homeostasis and oxidative stress management were strongly engaged, as indicated by the induction of superoxide dismutases, multiple glutathione S-transferases, catalase and other ROS-scavenging enzymes, together with auxiliary redox components such as glutaredoxin-S1-like and metallothionein-like proteins. The breadth and internal coherence of these redox-related functional groups support sustained management of oxidative load as stress progresses. This redox reinforcement was accompanied by a pronounced investment in proteostasis and selective protein turnover. Functional groups associated with protein folding, degradation and recycling were over-represented, including small heat shock proteins, regulatory subunits of the 26S proteasome, components of the ubiquitin–proteasome system, and the autophagy-related gene ATG10. Together, these changes indicate active removal of damaged proteins and maintenance of proteome integrity at late stages of salinity stress [78, 79].

Finally, GPE036890 showed a specific activation of a cell-surface remodeling program that is well aligned with acclimation to chronic salinity[80–82]. Functional categories related to cell wall organization, modification and biogenesis were prominently represented, as reflected by the regulation of cell-wall loosening and restructuring factors (Figure 6C - Figure 8C), including an expansin and the xyloglucan endotransglucosylase/hydrolase proteins XTH23 and XTH33, together with pectin-modifying enzymes such as pectin acetylesterase 11-like and pectinesterase 31. The coordinated modulation of these wall-related functional groups points to dynamic tuning of wall architecture and wall-derived signaling, with potential consequences for tissue mechanics, permeability and ion movement under sustained stress [80, 83]. Functional categories associated with extracellular barrier formation and lipid-based surface protection were engaged, as indicated by the induction of suberization- and cuticle-related components, including a suberization-associated anionic peroxidase 2 and a CER3-like wax synthase. This pattern is consistent with reinforcement of outer protective layers that may limit uncontrolled solute leakage and contribute to mitigation of oxidative injury during prolonged exposure [84].

Altogether, the amplitude and coherence of these genotype-specific changes suggests that tolerance in GPE036890 derives from network-level homeostatic tuning rather than from a limited set of “core” stress genes.

**Table 1.**
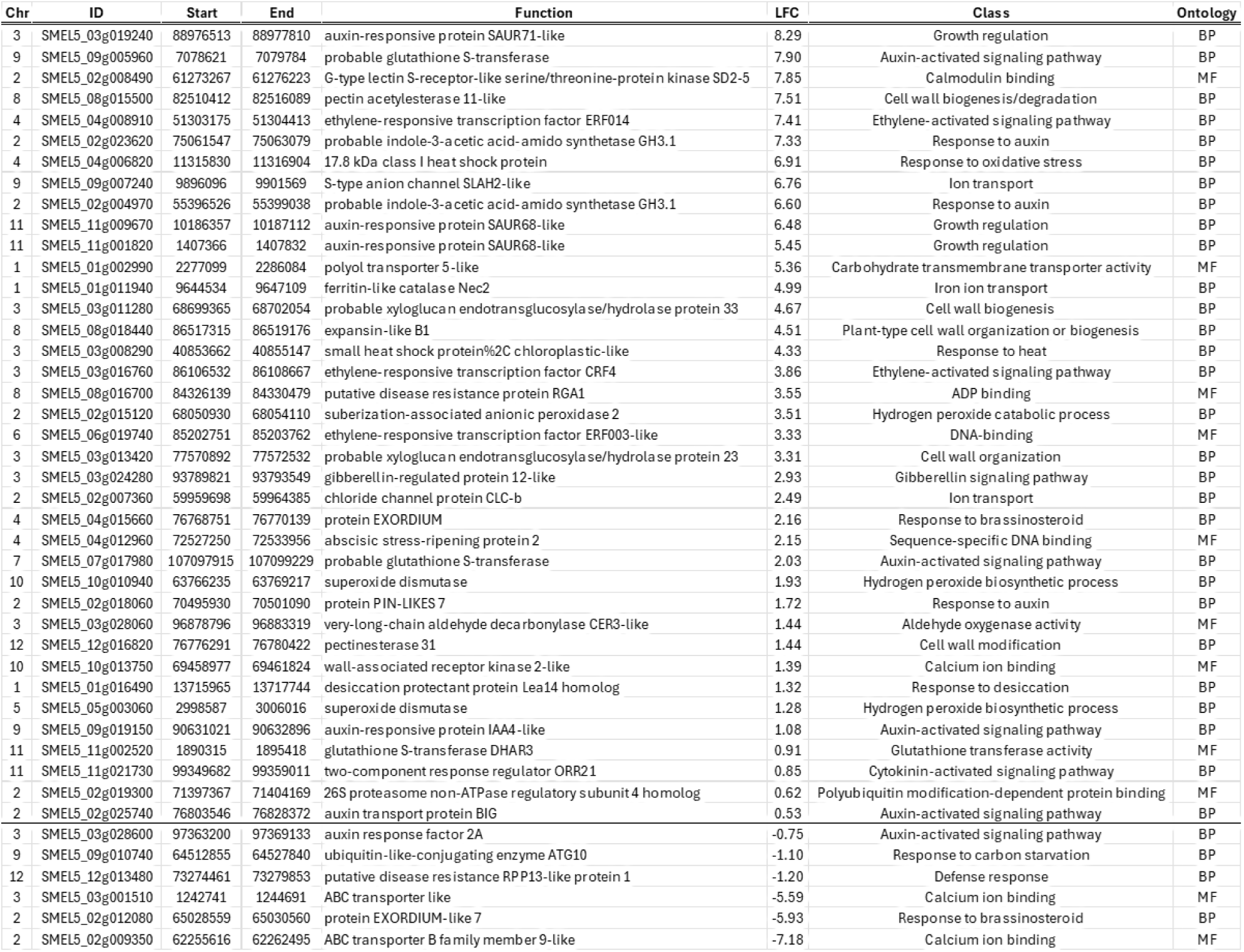
List of genes differentially expressed exclusively in the tolerant genotype (GPE036890) in the control vs salt comparison, with corresponding functional annotation, genomic coordinates (chromosome and start–end position), LFC, GO Class and Ontology. Only statistically significant DEGs (padj<0.05) are reported.

## Conclusions

From a breeding perspective, our results provide a hint on how combining multi-scale phenotyping with transcriptomics in a core-collection framework can disentangle global from genotype-specific determinants of salt tolerance. The conserved salt-stress transcriptional program we describe likely defines a necessary, but not sufficient, baseline, whereas the tolerant-specific reprogramming in GPE036890 suggests physiological strategies (maintenance of Ψ_leaf, cell-wall structure and K⁺/Na⁺ homeostasis) and associated candidate genes that are directly exploitable in breeding. A logical next step will be to develop segregating populations from crosses between GPE036890 and contrasting sensitive accessions, such as GPE022290, and to construct high-density genetic maps to resolve QTL for key physiological readouts, testing whether the candidate genes here identified co-localize with major-effect loci. In parallel, future work should couple such mapping efforts with rigorous physiological and multi-omics characterization across developmental stages and salinity levels, including gas exchange and chlorophyll fluorescence, quantitative ionomics of different tissues, and targeted metabolomic and hormone profiling in leaves and roots, to mechanistically link mapped loci and expression signatures to whole-plant salt adaptation.

## Author contributions

**M.M**: Writing – review & editing, Writing – original draft, Visualization, Methodology, Investigation, Formal analysis, Data curation, Conceptualization; **C.M**: Writing – review & editing, Writing – original draft, Visualization, Methodology, Investigation, Formal analysis, Data curation, Conceptualization; **A.M**: Writing – review & editing, Visualization, Conceptualization; **A.M.M**.: Writing – review & editing, Methodology, Formal analysis ; **L.B**: Writing – review & editing, Methodology, Investigation; **A.A**. : Writing – review & editing, Methodology, Investigation; **C.C**: Writing – review & editing, Methodology, Investigation, Conceptualization; **F.S**: Writing – review & editing, Methodology, Investigation, Conceptualization; **E.P**: Writing – review & editing, Methodology, Investigation, Conceptualization.

## Data availability

Sequencing data used in this study are openly available in the NCBI database.

---

**Supplementary Figure 1.**
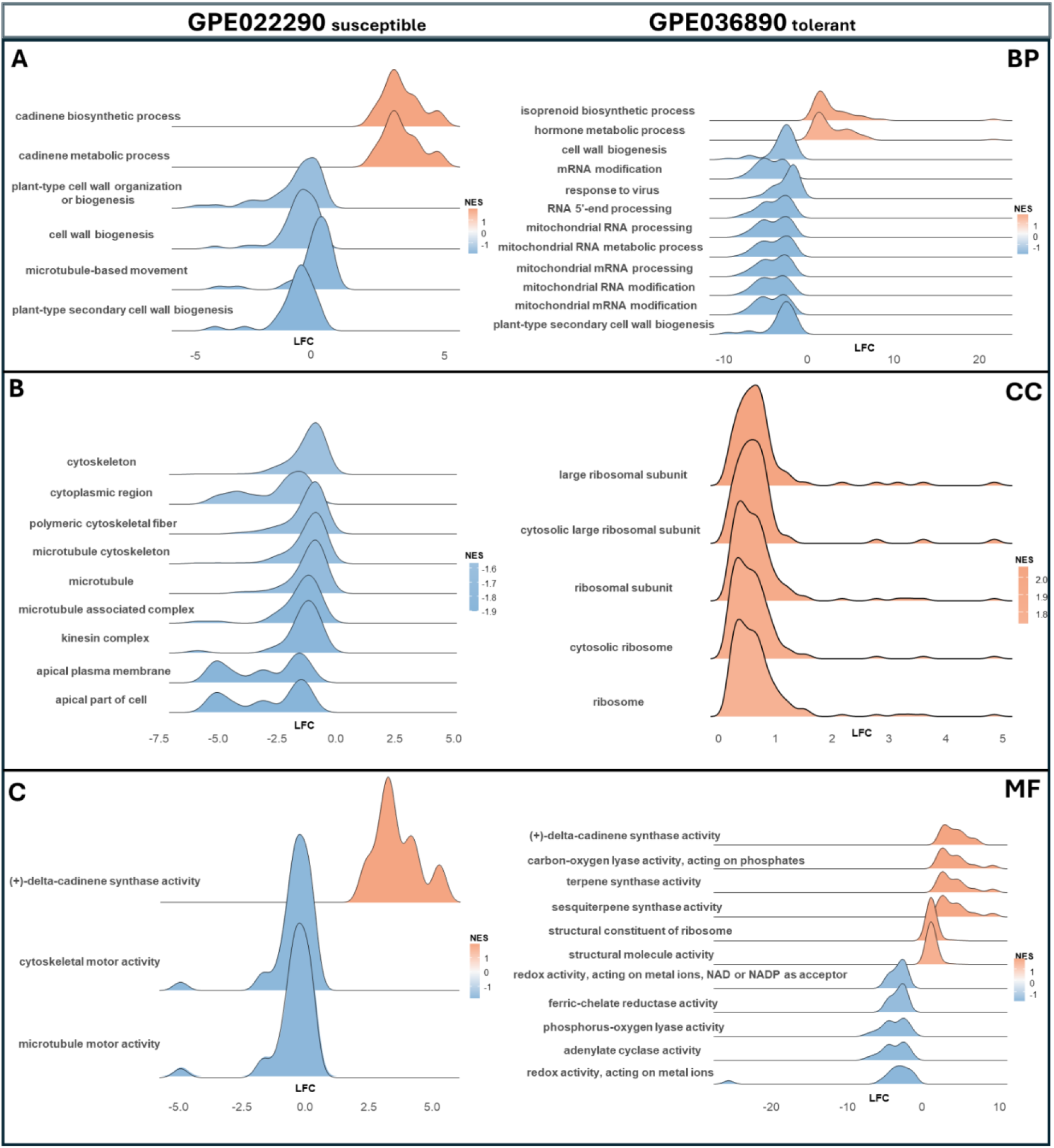
Gene set enrichment analysis (GSEA) of salt-induced transcriptional responses within each eggplant genotype. Ridge plots showing significantly enriched GO terms for biological process (A), cellular component (B) and molecular function (C) categories in the sensitive genotype GPE022290 and the tolerant genotype GPE036890 upon salt treatment, relative to their respective controls. The x-axis represents log₂ fold change (LFC), while the distribution shape reflects the contribution of individual genes to each term. Curves are filled with NES value, ranging from negative (blu) to positive (orange) values.

